# Stationary Equations for Non-Markovian Biochemical Systems

**DOI:** 10.1101/360123

**Authors:** Jiajun Zhang, Tianshou Zhou

## Abstract

We develop a new approach for stochastic analysis of biochemical reaction systems with arbitrary distributions of waiting times between reaction events. Specifically, we derive a stationary generalized chemical master equation for a non-Markovian reaction network. Importantly, this equation allows to transform the original non-Markovian problem into a Markovian one by introducing a mean reaction propensity function for every reaction in the network. Furthermore, we derive a stationary generalized linear noise approximation for the non-Markovian system, which is convenient to the direct estimation of the stationary noise in state variables. These derived equations can have broad applications, and exemplars of two representative non-Markovian models provide evidence of their applicability.

## Introduction

-The theory of Markov processes is well established, and has found its applications in an array of scientific fields including biology, chemistry, physics, epidemiology, ecology, and finance [1-3]. An important foundation of this theory is the chemical master equation (CME) [1,2,4], which can be simulated using numerical methods [5-8], or can be analytically solved in some cases [1,2,9-14].The mathematical tractability of Markov processes enables great simplifications in problem formulation, leading to important successes in the description of many stochastic processes ranging from gene regulation and mass transport to disease spreading and animal species interactions [1-3].

A Markovian biochemical process is memoryless, that is, the probabilities that future reaction events happen depend only on the present state of the system, independent of the prior history. However, many biochemical processes have memory or are non-Markovian. Non-exponentially distributed waiting times [15-18] and time delays [19-22] between reaction events can lead to memory. Non-Markovity has also been verified by the increasing availability of time-resolved data on different kinds of interactions [23-29]. The continuous time random walk (CRTW) provides a systematic starting point to account for arbitrary waiting time distributions between reaction events [30-38].

Recently, a generalized chemical master equation (gCME) was derived, which is capable of accounting for non-exponential interreaction times and the resulting non-Markovian character of reaction dynamics in time [38]. But the problem with the gCME is that the so-called “memory functions” are implicitly expressed by waiting time distributions and/or distributed delays. This leads to notorious difficulties in obtaining the system’s behavior, greatly limiting applications of the gCME (although numerical calculations can sometimes provide useful information [39-44]). While a dynamic distribution exactly captures stochastic behavior of a chemical reaction system, steady-state distribution is an important quantity needed to characterize the stationary behavior of the underlying system. To assess how experimental data can be informative, it is also needed to calculate or simulate aspects of steady-state distribution [45-47].

In order to obtain information on stationary behavior of a non-Markovian biochemical system, we introduce a mean reaction propensity function for every reaction to replace the memory function in the traditional treatment. Thus, the original ‘memory functions’ are currently converted into memoryless functions expressed explicitly by the integrals of the given waiting time distributions over the full time. Importantly, we derive a stationary gCME (sgCME), which allows to transform the original sticky non-Markovian problem into a mathematically tractable Markovian one. With this novel formulation, we further derive a stationary generalized linear noise approximation (sgLNA) for the original non-Markovian system, which allows the direct estimation of the stationary noise in state variables. These derived stationary equations can have broad applications and are particularly useful for the analytical derivation of stationary distributions in, e.g., a gene model of bursty expression with general waiting time distributions (Note: this is an issue unsolved in previous works).

### General Theory

-Consider a general chemical reaction network consisting of *N* different species (denoted by *X_j_*, *j* = 1, 2,…, *N*) that participate in *L* different reactions of the form

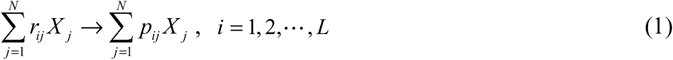

where *r_ij_* and *p_ij_* are stoichiometric coefficients, taking non-negative integers, and the differences *s_ij_* = *p_ji_* − *r_ji_* are stored in a *N* × *L* stoichiometric matrix **S**. Denote by *n_j_* the molecule number of species *X_j_* and by ***n*** = (*n*_1_,, … *n_N_*)^*T*^ the state vector, where T represents transpose. Let *ψ_i_* (*t*; ***n***) be the probability density function (PDF) of the *i*th reaction waiting time (depending on the system state, ***n***), and Ψ_*i*_ (*t*; ***n***) be the cumulative distribution function. In the following, we consider only the non-Markovity resulting from non-exponential waiting times between reaction events whereas the non-Markovity generated by distributed delays will be discussed later.

The time-evolutional gCME of the above network system has been established [31,33,34,38], but solving this equation has been thwarted to date. Instead, here we consider steady-state behavior of the system. Assume that the stationary probability of the system exists, and denote it by *P* (***n***). Then, based on the chemical CRTW theory [38], we can derive the following stationary equation (see the Supplemental Online Material [48] for its derivation)

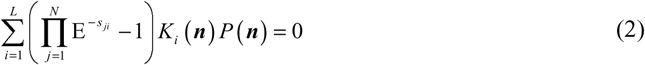

where *E*^−*s_ji_*^ is a step operator with the operation rule below: it removes *s_ji_* molecules from species *X_j_* in the *i*th reaction, i.e., *E*^−*s_ji_*^ *f* (***n***) = *f* (*n*_1_ ,…, *n_j_* − *s_ji_* ,…, *n_N_*) for any function *f* (***n***). Symbol *K_i_* (***n***) represents the mean propensity function of the *i*th reaction, accounting for the transition probability from a given state ***n*** to any other state. Importantly, we can show that *K_i_* (***n***) is explicitly expressed by given waiting time distributions, that is,

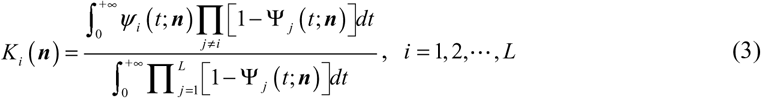

See the Supplemental Online Material [48] for the derivation of Eq. (3). Equation (2) and Eq. (3) altogether constitute the final sgCME, which governs stationary behavior of the original non-Markovian reaction system.

First, we observe that function *K_i_* (***n***) depends only on state variable ***n*** , independent of the prior history. In other words, the mean reaction propensity functions associated with the original non-Markovian process are memoryless. Then, we point out that the sgCME has important implications. For example, if we compare Eq. (2) with the common CME for the same structure reaction network with rate-limited reactions or with exponentially distributed waiting times (which is a certain Markovian chemical network), then we find that Eq. (2) is actually the stationary version of this CME. Moreover, if *K_i_* (***n***) is taken as the ‘reaction propensity function’ (possibly a rational rather than polynomial-type function of ***n***) of the *i*th reaction in this Markovian network, then we successfully convert a non-Markovian problem into a Markovian problem.

In particular, if the waiting times for some reaction are exponentially distributed, e.g., if *ψ* (*t*; ***n***) = *λ_i_* (***n***) *e*^−*λ_i_*(***n***)*t*^ I_(0,∞)_ (*t*) for some *i* , where I_*A*_ (*x*) is an indicator function of set *A* , i.e., I_*A*_ ( *x*) = 1 if *x* ∈ *A* and I_*A*_ ( *x*) = 0 otherwise, and *λ_i_* (***n***) represents the transition probability of this reaction, implying Ψ_*i*_ (*t*; ***n***) = 1 − *e*^−*λ_i_*(***n***)*t*^ I_(0,∞)_ (*t*) , then we find (see the Supplemental Online Material [48] for derivation)

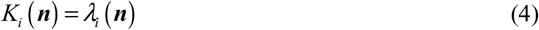

In other words, if the waiting times for some reaction follow an exponential distribution, then the corresponding *K_i_* (***n***) is equal to the reaction propensity function of this reaction. If all the reaction waiting times are exponentially distributed, then the corresponding sgCME is reduced to the common stationary CME (sCME).

To help the reader further understand the physical meaning of function *K_i_* (***n***) , we state some facts. First,
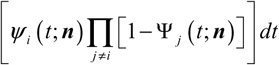
represents the probability that the *i*th reaction happens within the infinitesimal waiting time [*t*,*t* + *dt*], so the integral
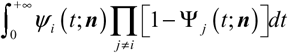
represents the cumulative possibility that the *i*th reaction takes place within the full time. Then, the equality
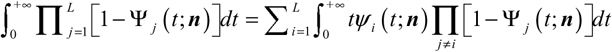
always holds, so the integral
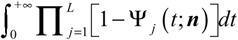
represents the mean waiting time that all the reactions happen. Thus, function *K_i_* (***n***) represents the mean transition probability of the *i*th reaction within the mean waiting time.

As an application of Eq. (2) with Eq. (3), we consider an interesting case where the waiting time for every reaction in the reaction network is assumed to follow a power-law distribution of Pareto type, that is, *ψ_i_* = *α_i_* (***n***) (*τ_i_*(***n***))^*α_i_*(***n***)^/*t*^*α_i_*(*n*)+1^ I_(0,∞)_ (*t*) [49]. This distribution is characterized by both a positive scale parameter *τ_i_* (***n***) and a positive shape parameter *α_i_* (***n***). Note that if *α*(***n***) ∈(0,1] , such a waiting time is a parsimonious model for infinite-mean random variables due to the generalized central limit theorem [49]. In this case, the corresponding system is weakly ergodicity breaking [50,51], which is a common characteristic of anomalous transport in heterogeneous environments and can lead to a fractional-in-time differential equation [50]. In spite of this, we can show (see the Supplemental Online Material [48] for detail)

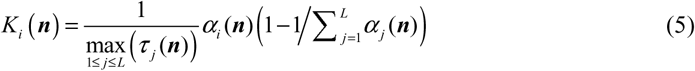

which is finite, independent of the convergence of raw moments of the power-law distribution.

Time delays, which account for the non-Markovian nature of many random processes, play a key role in many problems involving biochemical reactions or mass transport [19-22]. If a distributed delay is introduced to the above reaction network, then we can show (see the Supplemental Online Material [48] for derivation)

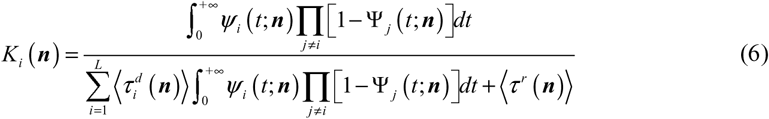

where
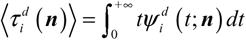
represents the mean delay time of the *i*th reaction with the PDF
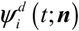
and is assumed to finite. If we denote by *τ^r^* the waiting time for the next reaction (i.e., the inter-reaction waiting time without delay), then
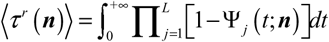
represents the mean inter-reaction waiting time. In particular, if
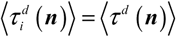
(which is called the mean global delay time) for 1 ≤ *i* ≤ *L* , i.e., if all the mean delay times
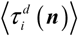
are equal, then we have (see the Supplemental Material [48] for derivation)

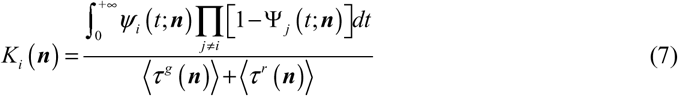

which indicates that the mean reaction propensity function of a reaction only depends on the mean global delay time, independent of the delay probability distribution. We point out that Eq. (7) is the generalization of a previous result in ref. [38] wherein an approximation was made.

Although we have derived the nice mathematical form of the sgCME for a general non-Markovian biochemical network (i.e., Eq. (2) with Eq. (3)), characterizing fluctuations in the individual reactive species is still difficult in some cases. On the other hand, it is well known that for a stochastic system, statistical quantities of the state variables, such as the first-order raw moments (or means) and the second-order central moments, are often most interesting since they can simply characterize the fluctuations in the state variables of the system. Here, we present a stationary generalized linear noise approximation (sgLNA) for the above non-Markovian system. Recall that for a given Markovian reaction system, the equations governing dynamics of the first-order raw moments (or means) and the second-order central moments of the state variables have been derived [1,2]. In order to calculate these statistical quantities in the above non-Markovian reaction system, we construct a Markovian biochemical process using *K_i_* (***n***). More precisely, we construct a Markovian reaction network such that it has the same structure as the original non-Markovian reaction network but takes *K_i_* (***n***) as the probability transition function of the *i*th reaction, where *i* = 1, 2,… , *L*. For such a constructed Markovian system, we can easily derive its rate equations, e.g., at steady state (see the Supplemental Online Material [48]), they are assumed to take the form [1,2]

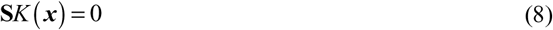

which is called a stationary generalized reaction rate equation, where *x* = ( *x*_1_, …, *x_N_*)^T^ with *x_i_* representing the concentration of reactive species *X_i_*, **S** = (*s_ij_*) is a stoichiometric matrix, and ***K*** (***x***) = (*K*_1_(***x***), … *K_N_* (***x***))^T^ is a *N*-dimensional vector. If the solution of Eq. (8) is denoted by ***x**_S_*, then ***x**_S_* is the vector of the stationary mean concentrations of the state variables in the original non-Markovian system.

In order to derive analytical formulae for calculating the second-order central moments of the state variables in the original non-Markovian reaction system, we adopt the Ω-expansion method [1,2]. Let Ω represent the volume of the system and write ***n*** = **Ω*x*** + **Ω**^1/2^ ***z***. For convenience, we introduce two matrices **A**_S_ and **D**_S_ , the entries of which are given by
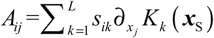
and
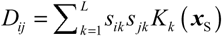
, respectively. Then, the covariance matrix ∑_S_ = (〈***x*** − ***x***_S_)(***x*** − ***x***_S_)^T^〉) satisfies the following Lyapunov matrix equation (see the Supplemental Online Material [48] for derivation)

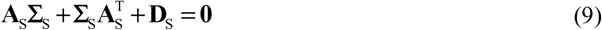

From this algebraic equation, we can easily obtain the stationary second-order central moments of the state variables in the original non-Markovian system.

### Implementations

-Here we apply the above general theory to two representative examples. First, birth-and-death processes, with some straightforward additions such as innovation, are a simple, natural and formal framework for modeling a vast variety of biological processes such as population dynamics, speciation, genome evolution [1,2,52]. Therefore, the first example we will analyze is a generalized birth-death process, which constitutes a fundamental model of non-Markovian evolutionary dynamics. This process can be described by two reactions:
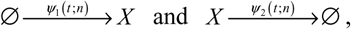
where *ψ*_1_ (*t*; *n*) and *ψ*_2_ (*t*; *n*) are waiting time distributions of birth and death respectively, and *n* represents the molecule number of species *X*. This model is general and can include almost previously studied models of birth-death processes as its special cases [1,2,52]. According to the above sgCME, the steady-state equation corresponding to this process is given by [48]

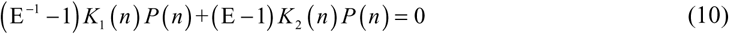

where *K_i_* (*n*) (*i* = 1, 2) can be obtained through Eq. (9). From Eq. (10), we can obtain the iterative relation: *K*_2_ (*n*) *P*(*n*) = *K*_1_ (*n* − 1) *P* (*n* − 1) , from which we can further obtain the explicit expression of stationary distribution

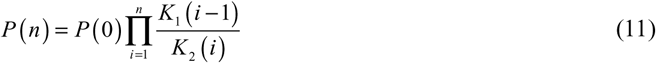

In particular, if *ψ*_1_ (*t*, *n*) = *ψ*_1_ (*t*) I_(0,∞)_ (*t*) , and *λ*_2_ (*t*, *n*) = *λ*_2_ *ne*^−*λ*_2_*nt*^ I_(0,∞)_ (*t*) , then we can have

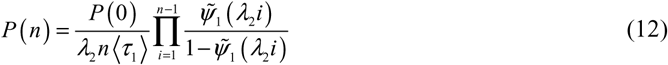

where *ψ̃*_1_ (·) is the Laplace transform of function *ψ*_1_ (*t*) , and
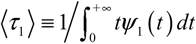
is the reciprocal of the mean birth waiting time. Furthermore, if
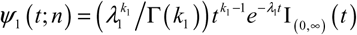
in which Γ(·) is the common gamma function, then we can obtain stationary distribution *P* (*n*) = ( _0_*F*_1_ (; 2*λ*; *λ*^2^))^−1^ *λ*^2*n*^/((2*λ*)_*n*_ *n*!) , where we denote *λ* = *λ*_1_/*λ*_2_ and _0_*F*_1_ is a generalized hypergeometric function [53]. In addition, we can obtain the analytical mean and variance given by 〈*n*〉 = (2^1/*k*_1_^ − 1)*λ* and ∑ = 2^1/*k*_1_−1^ *λ*/*k*_1_ respectively. The Fano factor, which is defined as the ratio of the variance over the mean, is given by *F* = 2^1/*k*_1_−1^/[(2^1/*k*_1_^ − 1) *k*_1_. In the Supplemental Material [48], we also analyzed the case that waiting times for the birth process follow an exponential distribution and those for the death process follow a general distribution. In addition, we presented a numerical method for the case that waiting time distributions for birth and death processes are all general.

Numerical results are demonstrated in Fig. 1. From this figure, we observe that results obtained by sgCME and by sgLNA are in good agreement. Similarly, results obtained by Gillespie stochastic simulation (lines in Fig. 1(a)) are also in good accord with those obtained by theoretical prediction (empty circles in Fig. 1(a)). In addition, we observe from Fig. 1(b) and 1(c) that the larger the *k*_1_ is, the smaller are the mean and the Fano factor, implying that non-Markovity can reduce the mean and noise of the outcome. Note that *k*_1_ = 1 corresponds to the case of exponential waiting times.

**Fig. 1.**
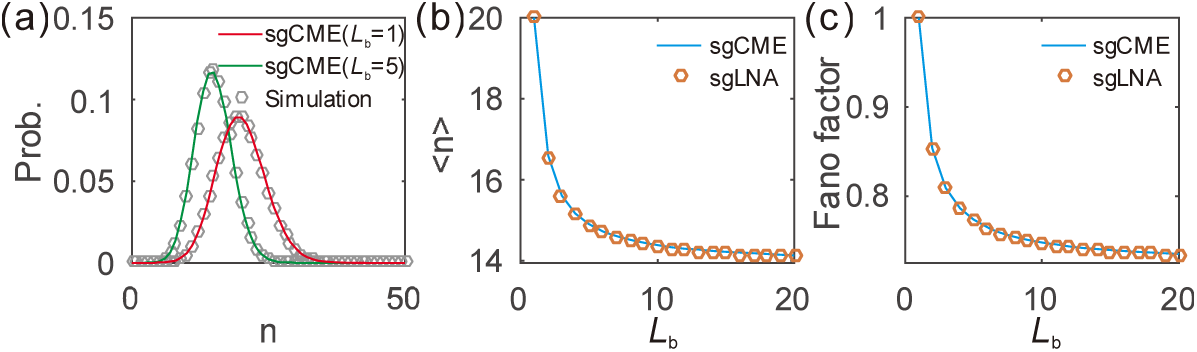
Comparison between results obtained by sgCME, and those obtained by sgLNA (seeing the analytical results below Eq. (12)), in a non-Markovian birth-death process, (a) Probability distribution; (b) Dependence of mean on *L_b_* (= *k*_1_); (c) Dependence of Fano factor on *L_b_*. Some parameter values are set as *k*_1_ = *L_b_* , *λ*_1_ = 10*L_b_* , *λ*_2_ = 1 , where either the values of *L_b_* are indicated in (a) or its change range is set in (b) and (c).

As a simplification, the dynamics of gene expression probabilities is described often by coupled birth-death processes, where birth corresponds to protein synthesis while death occurs via degradation. Therefore, we next consider a model of gene self-regulation, which is described by
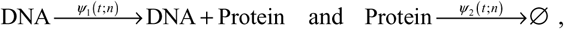
where *n* represents the number of protein molecules [11,54,55]. We point out that this model belongs to the above discussed case in the absence of feedback, but it can be a more general birth-death process in other cases. For analytical consideration, we assume that waiting time distributions for protein birth and death take the forms: *ψ*_1_ (*t*; *n*) = ((*λ*_1_(*n*))^*k*_1_^/Γ(*k*_1_))*t*^*k*_1_−1^*e*^−*λ*_1_(*n*)*t*^ I_(0,∞)_ (*t*) where *λ*_1_(*n*) is a function of *n* , and *ψ*_2_ (*t*; *n*) = *λ*_2_ *e*^−*λ*_2_*t*^ I_(1,∞)_ (*t*) , respectively. Furthermore, we consider the case that function *λ*_1_(*n*) is linear in *n* , e.g., *λ*_1_ (*n*) = (*n* + *n*_0_)*λ*_1_ corresponding to linear feedback) is set, where *λ*_1_ is a constant and *n*_0_ is a positive integer. By simple calculations, we can show *k*_1_ (*n*) = [(*nλ*_2_ (*n*+*n*_0_)*λ*_1_)^*k*_1_^]/[(*nλ*_2_ + (*n* + *n*_0_)*λ*_1_)^*k*_1_^ − ((*n* + *n*_0_)*λ*_1_)^*k*_1_^] and *K*_2_(*n*) *nλ*_2_. In particular, if *k*_1_ = 1, *n*_0_ = 1, then *P* (*n*) = (1 − *ρ*)^*n*^ *ρ* is a geometric distribution, where *ρ* = 1 − *λ*_1_/*λ*_2_ and *n* = 0,1, 2,…. For the case of nonlinear feedback, similar analysis can be carried out but the results are more complex.

In contrast to the above analyzed gene model that corresponds to constitutive expression, there is another expression way, i.e., bursty gene expression. Therefore, we next consider the second example, which is an on-off model of bursty gene expression with non-exponential waiting times (referring to Fig. 2(a)). The biochemical reactions be described by four reactions:
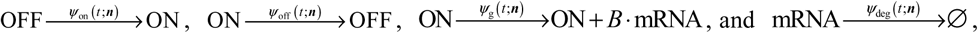
where functions *ψ*_on_ (*t*; ***n***) , *ψ*_off_ (*t*; ***n***) , *ψ*_g_ (*t*; ***n***) and *ψ*_deg_ (*t*; ***n***) are all waiting time distributions, state vector ***n*** = (*n*_1_ , *n*_2_ , *m*)^*T*^ with *n*_1_ = *off* and *n*_2_ = *on* as well as *m* representing the numbers of DNA molecules at off and on states as well as the number of mRNA molecules respectively, and the burst sizes *B* follow a geometric distribution *P_b_* ( *B* = *k*) = *b^k^*/(1 + *b*)^*k*+1^ (*k* = 0,1,…) with *b* being the average burst size. For analytical consideration, we assume that on and off waiting times follow respectively Erlang distributions *ψ*_on_ (*t*; ***n***) = ((*λ*_on_ *n*_1_ *f*)^*k*_on_^/Γ(*k*_on_))*t*^*k*_on_−1^*e*^−*λ*_on_*n*_1_*t*^ and *ψ*_off_ (*t*; ***n***) = ((*λ*_off_*n*_2_)^*k*_off_^/Γ(*K*_off_))*t*^*k*_off_−1^*e*^−*λ*_off_*n*_2_*t*^, and transcription and degradation waiting times follow respectively exponential distributions *ψ*_g_(*t*; ***n***) = *μn*_2_ *e*^−*μn*_2_*t*^ and *ψ*_deg_ (*t*; ***n***) = *λ*_deg_ *me*^−*λ*_deg_*mt*^. Then, we can derive the analytical expressions of *K_i_* (***n***) (*i* = 1, 2,3, 4) (see the Supplemental Online Material [48] for detail). Numerical results are demonstrated in Fig. 2(b-d).

**Fig. 2.**
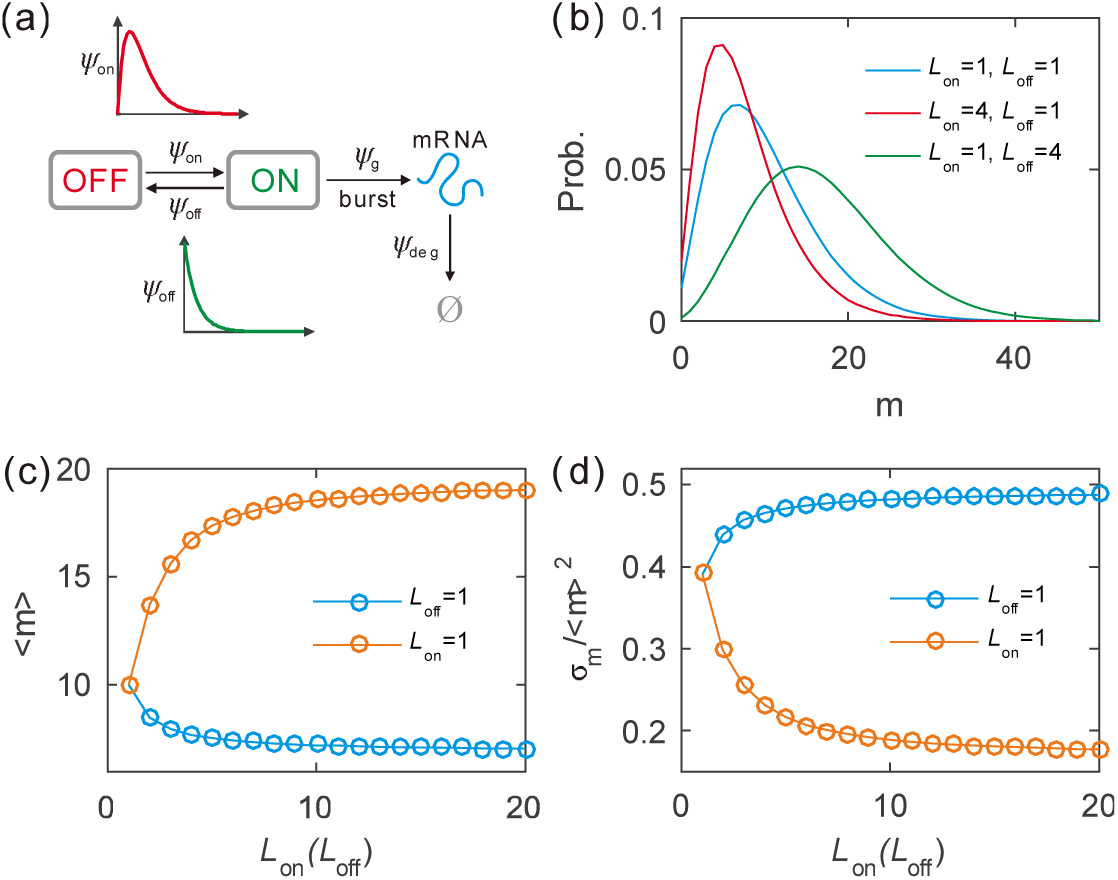
Effect of non-Markovianity on gene expression. (a) Schematic for a model of stochastic transcription, where the times that the gene dwells at ON and OFF states follow general distributions, and mRNA is synthesized in a bursty manner and degrades in a multistep manner (leading to a non-exponential waiting-time distribution). (b) Stationary mRNA distributions in three representative cases. (c) Dependence of the mRNA mean on *L*_on_ if *L*_off_ is fixed or on *L*_off_ if *L*_on_ is fixed. (d) Dependence of the mRNA noise intensity on parameter *L*_on_ (or *L*_off_). Some parameter values are set as *μ* = 10 and *b* = 2 , whereas other parameters are determined by relations: *k*_on_ = *L*_on_, *λ*_on_ = 5*L*_on_, *k*_off_ = *L*_off_ and *λ*_off_ = 5*L*_off_.

From Fig. 2(c), we observe that if *L*_on_ = 1 is fixed, the mRNA mean is monotonically increasing in *L*_off_ but if *L*_off_ = 1 is fixed, it is monotonically decreasing in *L*_on_. In both cases, the mRNA mean almost keep unchanged after a certain value of *L*_on_ or *L*_off_. However, the change tendency for the rate of the mRNA variance over the square of the mRNA mean is opposite to that of the mRNA mean (comparing Fig. 2(d) with Fig. 2(c)). Figure 2(b) shows stationary mRNA distributions in three special cases, which can further verify the change tendency in Fig. 2(c) and 2(d). These analyses indicate that non-Markovity plays an unneglectable role in affecting gene expression.

### Conclusions

-We have derived an exact sgCME and a sgLNA from the gCME for a general reaction network with arbitrary (exponential or non-exponential) waiting time distributions or/and with distributed delays. These derived equations allow one to retain analytical and/or numerical tractability, being general in scope, and thus of a potential applicability in a wide variety of problems that transcend pure physics applications. The derived sgCME is particularly useful in deriving stationary distributions in some sticky non-Markovian biochemical systems, as demonstrated in this article. The power of the sgCME can be enhanced by analyzing other examples such as non-Markovian random walks and diffusion on networks [56-63], and non-Markovian open quantum systems [64]. We expect that our analytical frameworks will be of use for studies of a variety of phenomena in biological and physical sciences, and indeed in other areas where individual-based models with general waiting time distributions and/or delayed interactions are relevant.

This work was supported by grants 91530320, 11775314, 11475273, and 11631005 from Natural Science Foundation of P. R. China; 2014CB964703 from Science and Technology Department, P. R. China; 201707010117 from the Science and Technology Program of Guangzhou, P. R. China.

